# Substitutional landscape of a split fluorescent protein fragment using high-density peptide microarrays

**DOI:** 10.1101/2020.05.20.105668

**Authors:** Oana N. Antonescu, Andreas Rasmussen, Nicole A.M. Damm, Ditte F. Heidemann, Roman Popov, Alexander Nesterov-Mueller, Kristoffer E. Johansson, Jakob R. Winther

## Abstract

Split fluorescent proteins have wide applicability as biosensors for protein-protein interactions, genetically encoded tags for protein detection and localization, as well as fusion partners in super-resolution microscopy. We have established and validated a novel platform for functional analysis of leave-one-out split fluorescent proteins (LOO-FPs) in high throughput and with rapid turnover. We have screened more than 12,000 strand 10 variants using high-density peptide microarrays for binding and functional complementation in Green Fluorescent Protein. We studied the effect of peptide length and the effect of different linkers to the solid support and mapped the effect of all possible amino acid substitutions on each position as well as in the context of some single and double amino acid substitutions. As all peptides were tested in 12 duplicates, the analysis rests on a firm statistical basis allowing determination of robustness and precision of the method. We showed that the microarray fluorescence correlated with the affinity in solution between the LOO-FP and peptides. A double substitution yielded a peptide with 9-fold higher affinity than the starting peptide.

## INTRODUCTION

Tens of new split fluorescent proteins (splitFPs) have been developed since the first reassembly of a splitFP was achieved 20 years ago (1). Fluorescent proteins (FPs) have been split in many creative ways, by removing fragments ranging from around half of the FP β-barrel (1) to one (2) or two (3) secondary elements. The splitFP fragments obtained have low or no fluorescence on their own, but can reassemble to form a fully functional FP. These properties have made splitFPs desirable for many bioanalytical applications, from sensing protein-protein interactions to protein detection and localization, and as tools in super-resolution microscopy (4, 5).

Leave-one-out splitFPs (LOO-FPs) are variants of splitFPs in which one of the secondary elements, such as one of the β-strands or the internal α-helix, typically of less than 20 amino acids, are removed (6). Ideally, left-out elements can spontaneously associate with the LOO-FP to recover fluorescence, making them useful as tags fused to a protein of interest, as well as individual peptides for *in vitro* applications. Preferably, a peptide should have high solubility, affinity and brightness upon complementation of the LOO fragment, as well as a small size that interferes minimally with a potential fusion partner protein (2, 5, 7, 8). Amino-acid substitutions in the sequence of the LOO-FP fragments offer the possibility to modulate their complementation efficiency, spectral properties, solubility and photostability. Although these are clear targets for optimizing LOO-FPs, genetic engineering methods such as random mutagenesis are laborious and do not directly measure binding between the splitFP fragments (9).

High-density peptide microarrays provide a powerful technology for massively parallel screening of peptide-protein interactions. While DNA and RNA arrays have been extensively used in mappings of polynucleotide-protein interactions (10), peptide microarrays have mostly been reserved for antibody epitope mapping and screening receptor-ligand interactions (11). Peptide microarrays cannot, however, typically accommodate peptides longer than 15-20 residues and, in addition, the affinity of the peptide-protein interaction to be investigated needs to be less than μM (11). Thus, peptide microarray analysis is attractive for complementing LOO-FPs, as peptides are generally under 20 residues (6), and can bind the LOO partner with dissociation constants from hundreds of picomolar to hundreds of nanomolar (4).

LOO-FPs systems can be divided in those where the chromophore is matured prior to reconstitution of the full-length protein and those were maturation takes place on reconstitution. While fluorescence recovery in the former is rapid (in the order of minutes) and essentially dependent on the rate of association of the partners, the latter may take hours requiring chemical condensation and oxidation of the chromophore. We hypothesized that, since binding of the split fragments with a preformed chromophore generates a fluorescent signal, their association could directly be followed by fluorescence detection on a peptide microarray. We tested this hypothesis using the *superfolder* “GFP split10” system, in which strand 10 is removed from the N-terminus of a circular permutated variant by trypsin digestion generating LOO10-GFP (12). The peptide microarray that we tested had a library of 12,544 left-out strand 10 (s10) peptide sequences in 12-fold repeats and was screened for complementation to the partner truncated protein using a fluorescence laser scanner. Contrary to DNA-based screening methods, chemical synthesis of defined peptides allowed for highly targeted analysis in which specific sequences can be queried in a non-random fashion with direct, rapid and quantitative readout.

We generated comprehensive splitFP sequence-function maps in a single experiment and with high precision, without the need of mutant selection rounds, enrichments or individual handling of clones. Analyzing peptides scanned with all possible amino acid substitutions in s10, we mapped hotspot residues and discovered improving substitutions. SplitFP complementation using peptide arrays also provided information about the sequences with lower fluorescence yield, which are generally inaccessible by genetic methods (2, 7). By introducing variations in the s10 context such as the peptide length and charge of the C-terminal array surface linker, we demonstrated the robustness of the assay. Finally, we assessed the accuracy of the microarray platform by characterizing interesting s10 sequences spectrally and thermodynamically in solution.

## RESULTS

### Experimental setup

LOO10-GFP was obtained from a circularly permuted *superfolder* GFP (cp-sfGFP), with β-strand 10 engineered in the N-terminal, employing minor modifications to an established protocol (12). S10 was removed by trypsin digestion followed by size exclusion chromatography in denaturing conditions, to yield LOO10-GFP (**Figure 1A**). Upon refolding in native buffer, LOO10-GFP was obtained in a fairly stable and soluble state which recovered fluorescence when reassembled with synthetic s10 variants in solution (data not shown).

**Figure 1.**
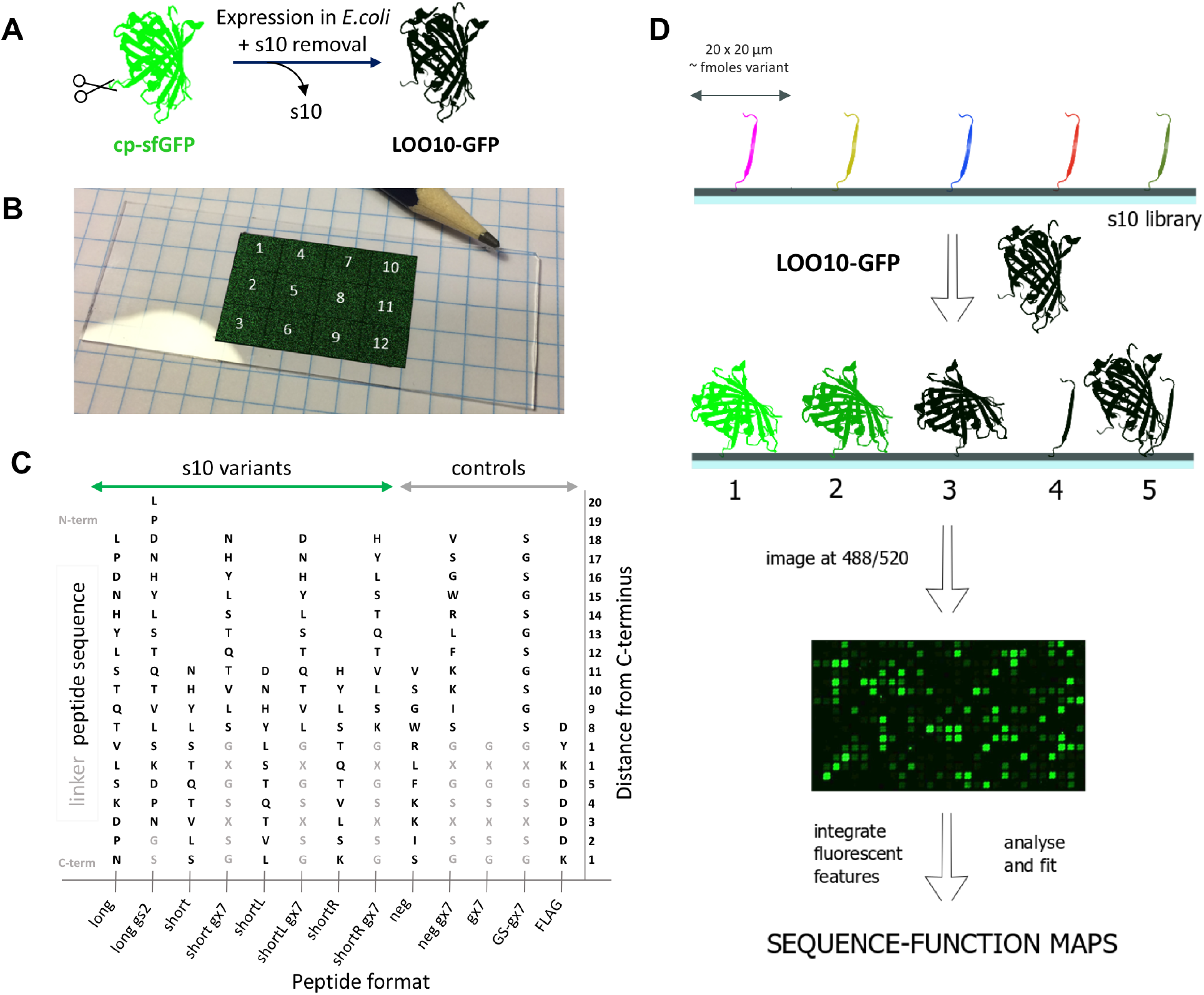
Experimental setup. (A) LOO10-GFP is obtained by removing strand 10 from a circularly permuted superfolder GFP; (B) Peptide microarray layout with 12 identical sectors (pencil tip for size appreciation). In each sector, peptide variants are bound to the solid surface at the C-terminus, forming well defined rectangular 20 x 20 μm spots - here pasted onto the slide in false color for visualization. (C) The peptide formats present in the library are long and short length variants with or without a gs2 or gx7 linker. Controls are also synthesized with or without linker. Linker sequences tested are gs2 – GS and gx7, which can be gs7 – GSGSGSG, gk7 – GKGSKSG or ge7 – GEGSESG. S10 and control sequences are colored black, while the linker sequences light gray. Substitutions were only performed in the s10 sequences and not in the linkers. (D) The peptide microarray is incubated with LOO10-GFP and imaged in the green channel. Scenarios at the microarray surface upon incubation: 1. Specific binding and high fluorescence; 2. Specific binding and low fluorescence; 3. Specific binding and no fluorescence; 4. No binding; 5. Non-specific binding. The fluorescence images were registered to the sequence information, then integrated and fit to sequence-function maps.

The s10 peptide library was designed as a single microarray layout with 12 identical sectors, amounting to 150,528 peptide spots in total on a microscope-format slide (shown in the false-color image **Figure 1B**). The s10 library is developed around the 18-mer “wild-type” s10 peptide L195PDNHYLSTQTVLSKDPN212 (termed s10_long_). Preliminary experiments had shown that a truncation to a core sequence of 11 residues, N_198_HYLSTQTVLS_208_ (s10_short_), forming a minimal β-strand, was also able to complement in solution (data not shown). In addition, preliminary microarray screens showed that 7-mer charged linkers for the short format would be beneficial (data not shown). Thus, 7-mer linkers with positive, neutral and negative charges were screened for s10_short_ in order to increase signal to noise, but also to assess signal robustness. We also wished to study how 1 amino acid shifts to the left and right of the sequence of s10_short_ influenced fluorescence recovery, resulting in the s10_shortL_ D197-L207 and the s10_shortR_ H199-K209 variants, respectively. An overview of the length, linker and substitution variants tested for the s10 peptide can be found in **Figure 1C** and **Table S1**. To generate a comprehensive picture of the sequence tolerance, all single amino acid substitutions were synthesized for the s10_long_ (342 variants), s10_short_ (209 variants), s10_short_L (209 variants) and s10_short_R (209 variants) formats. In addition, a series of double (1607 variants for s10_long_, 942 variants for s10_short_) and triple substitutions (304 variants for s10_long_, 171 variants for s10_short_) were added at positions 199, 203 and 207. Particular substitutions at positions 199 and 203 had in preliminary screens shown enhanced fluorescence, while substitutions at positions 207, were included in libraries as negative controls. To access the effect of linkers, all the single, double and triple substitutions were assessed both without linker and linked to GS (*gs2*) for s10_long_ and GSGSGSG (*gs7*), GKGSKSG (*gk7*) and GEGSESG (*ge7*) for s10_short_. As controls, substitutional scans of a negative control (*neg*) 11-mer peptide inspired from a split-luciferase (13) were synthesized to test the specificity of LOO10-GFP to s10 variants. We furthermore added linker and FLAG synthesis controls, and blank spots for background estimation.

Arrays were synthesized on modified surface microscope slides using a lithographic method described previously (14) and were purchased from a commercial vendor (Schafer-N, Copenhagen). Identical peptides were placed at the same position relative to other peptides within individual sectors, but peptides were deliberately *not* grouped by linker or composition within a sector, but rather placed randomly to avoid local artifacts affecting similar peptides in a similar fashion.

To determine binding and recovery of activity, we incubated the peptide microarray with LOO10-GFP, and after a 10 min wash step the microarray was briefly dried and imaged at 488 nm excitation and 520 nm emission using a microarray scanner (see Methods for details; **Figure 1D**). Image analysis of the fluorescent spots allowed us to assign each peptide in the library a fluorescent signal.

### Microarray assay performance

Before determining any substitutional effects, we quantitatively assessed the performance of the microarray assay in terms of precision, specificity and robustness. Data handling and statistical analyses are detailed in Methods. The high precision of the microarray method was proven by correlating the variant libraries in all 12 sector replicas, which showed Pearson coefficients > 0.94 (**Figure S1**).

Binding to positively charged peptides by low pI proteins is a common concern with peptide microarrays (15, 16). We tested whether the low-fluorescence LOO10-GFP might have nonspecific bias towards highly charged peptide spots, leading to false-positive signals. Results showed that microarray fluorescence was independent of formal peptide charges and distributes around s10_long_ WT, which has a formal net charge of −1 (**Figure 2A**). In addition, the fluorescent complementation was specific towards s10 variants, signal from all negative control peptides being minimal (**Figure 2B**).

**Figure 2.**
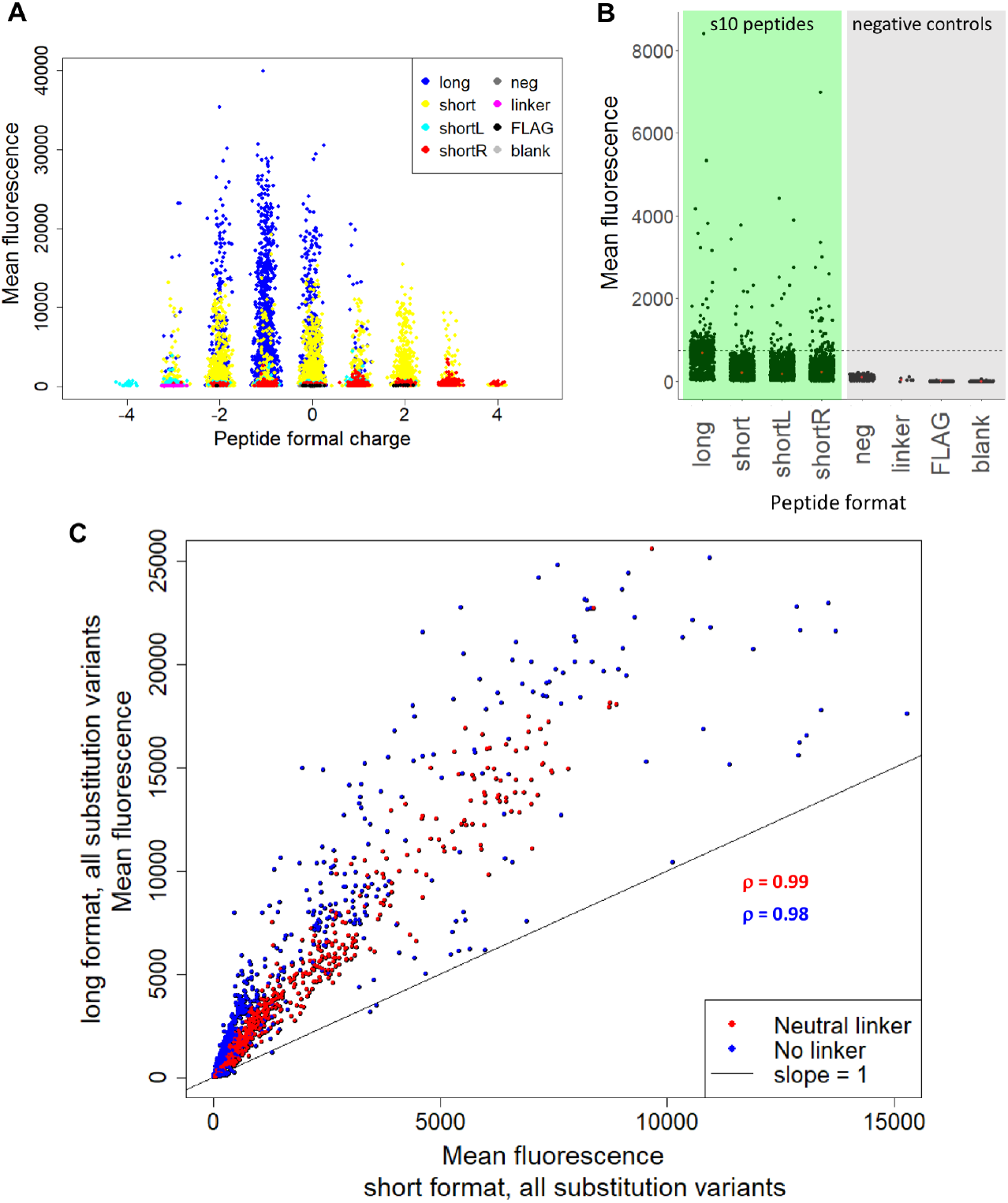
Microarray assay performance. (A) Mean fluorescence against formal charge of each peptide variant. The charge was estimated by counting all positive (R, K) and negative (D, E) charges in each sequence. (B) Mean fluorescence of linker and substitution variants for the long, short, shortL and shortR s10 peptides (green background), compared to neg, linker, FLAG and blank controls (grey background). The dashed line shows the fluorescence of s10_long_ WT, that we used as positive control in this dataset. Double and triple substitutions were excluded, since they were only performed on the long and short formats. (C) Red points indicate mean fluorescence of all variants with neutral linker in the short format versus the same variants of long format. Blue points indicate mean fluorescence of all substitution variants without linker of the short format versus the same variants of long format. The indicated p values are Pearson coefficients of the two correlations.

Comparing long and short length variants we found that peptide length and charged C-terminal linker variants can have an influence on the dynamic range of fluorescence. In **Figure 2C** we compared the same substitution variants in the long versus the short peptide format, with or without a neutral linker. The long and short variants correlated well with each other (Pearson coefficients > 0.98), but the signal from s10_long_ peptides was substantially higher than from s10_short_ format in terms of the dynamic range.

The various short formats are all well correlated for individual substitutions among themselves (**Figure S2**). For the s10_short_ format a linker is clearly preferable, presumably to increase the distance to the microarray surface. The shifted variants had low signal-to-noise, but seem to follow a nearly one-to-one relationship with short variants (**Figure S3**). This suggests that the 9-mer central sequence HYLSTQTVL could potentially work *per se* as a split 10 fragment.

The main conclusion from this comparison is that although there are differences in the performance of individual linker designs, the variant sequences correlated very well with each other between linker formats. Overall, the microarray signals are highly robust with regards to ranking across substitution variants. However, due to the different dynamic ranges obtained at different length and linker formats, substitutional effects should always be interpreted within the same peptide format.

### Effect of substitutions on s10_long_ array signal

Since all peptide formats gave sequence-dependent correlated signals, we will in the following discuss substitutional effects in the s10_long_ format without linker, since this format yielded the highest signals and dynamic range. The heatmap in **Figure 3A** shows the effect of all s10_long_ single residue substitutions on the fluorescence of the s10_long_:LOO10-GFP complex. Some increase in fluorescence when substituting T203 with hydrophobic residues Y, F, I and V was expected, because these substitutions are known to cause a red shift in the fluorescence emission maximum from 506 nm to, depending on the context, 515-527 nm (12, 17–19). The readout using a 520 nm emission filter could favor the T203 red-shifted substitutions. Besides the effects at position 203, replacing H199 with a series of hydrophobic residues (Y, F, I, V, L) also resulted in increased signal. Specifically, introducing H199Y as a single substitution offered 10-fold greater fluorescence compared to WT.

**Figure 3.**
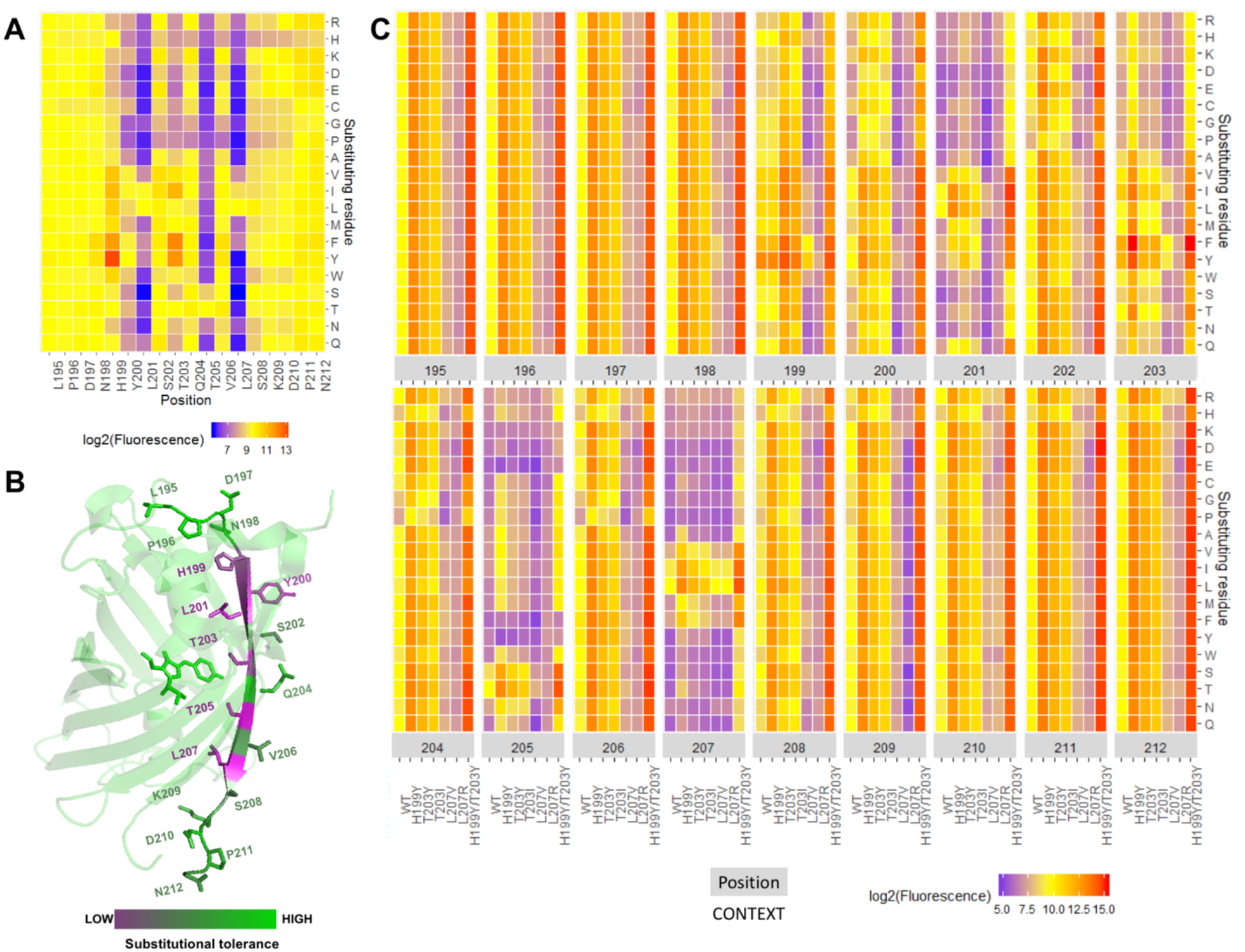
Effect of substitutions on s10_long_ fluorescent signal. (A) Saturation substitution heatmap of the s10_long_ peptide. (B) Substitutional tolerance values from 1 (low) to 19 (high) are plotted in a magenta-green gradient on the superfolder GFP with the s10 side chains and chromophore represented as sticks and LOOI0-GFP as cartoon (PDB: 2B3P). (C) Fluorescence heatmap of single, double and triple substitutions from s10 WT. Fluorescence of substitutions variants (y-axis) at each sI0 position (higher x-axis) in the WT, H199Y, T203Y, T203I, L207V, L207R and H199Y/L203Y context (lower x-axis). Median relative standard deviation RSD% across all variants was 15%. Color key of A and C heatmaps: Yellow – WT-like fluorescence; Blue – loss-of-function substitutions; Red – gain-of-function substitutions.

To evaluate the importance of each of the WT residues, we assigned each s10 position a substitutional tolerance value from 1 (lowest) to 19 (highest) based on the number of substitutions it could accept at each position without gaining or losing function, as described in Methods. Mapping this onto the superfolder GFP crystal structure (**Figure 3B**) showed that positions with lowest substitutional tolerance were 199, 200, 201, 203, 205, 207, all pointing inside the FP β-barrel except position 200. The low tolerance positions delimit the 199-207 region as the s10 core residues for function. Within core region, changing outward-pointing residues 202, 204, 206 to β-sheet-disruptive P and G were, as expected, detrimental (20). We noted that substitution to H had a negative impact on almost all positions.

In the peptide library we included full substitutional scans of s10_long_ H199Y, T203Y, T203I, L207V, L207R and H199Y/T203Y peptides, these libraries representing double and triple substitutions from WT. A full overview of the functional recovery of all tested single, double and triple substitutions from s10 WT is shown in the heatmap in **Figure 3C**. Most of the substitutional variants in the H199Y, T203Y and T203I contexts were distributed at fluorescence values above WT, increasing the fluorescence on average by a factor of 9.1, 4.2 and 2.4, respectively, but there were also many with less signal than WT (**Figure S4**). In the H199Y/T203Y context, which increased the fluorescence on average by a factor of 23, almost all substitutions are gain-of-function, with some exceptions for substitutions in positions 205 and 207. On the other hand, all substitutions in the L207R context are loss of function, meaning that there is no substitution that can rescue L207R. L207V context peptides are also distributed below WT, but there are some rescue substitutions in positions 199 and 203.

H199Y was the most fluorescence enhancing single substitution and the most robust when combined with other substitutions, obvious from both **Figure 3C** and **Figure S4**. Combining enhancing substitutions at positions 199 and 203 was advantageous. The best performing s10 peptide in the library, H199Y/T203F, with a 54-fold increase in fluorescence compared to WT, is closely followed by H199Y/T203Y combined with P211D or N212 to A, K, D, R and I, respectively. (**Figure S5A**). The many advantageous substitutions to charged residues in the C-terminal flanking region outside the hotspot could potentially suggest a need for increased peptide charge. In the core region, additional beneficial substitutions include L201I and V206T which are confirmed in the absence of flanking regions in the s10_short_ format (**Figure S5B**).

We found that most of the detrimental substitutions observed in the WT context are rescued above WT levels in the H199Y and T203Y contexts, except some substitutions in positions 201, 205 and 207 (**Figure 3C**). For example, L207V is rescued at 3-fold WT levels in presence of H199Y, while L207R is not. Only substitutions in positions 201, 205 and 207 can bring the fluorescence of variants in the high-performing H199Y/203Y context below WT levels (**Figure S4**), suggesting that these are the three most constrained s10 positions.

### How well does array signal reflect binding affinity in solution?

The effect of the most beneficial substitutions was hypothesized to reflect an affinity increase at sub-saturation concentrations of LOO10-GFP (**Figure S6**). On the other hand, intrinsic brightness of FP complexes at the wavelength of measurement would also affect apparent intensity. To address this issue, we chose to analyze the binding affinity and brightness, in solution, of the reconstituted GFP with the s10 WT peptide and three gain-of-function variants (H199Y, T203Y and H199Y/T203Y) in the short format and one loss-of-function variant in the long format (L207R). We examined whether their microarray signals correlated with brightness and/or affinity of the peptide variants upon reassembly with LOO10-GFP in solution.

Upon complementation of LOO10-GFP at saturating concentrations of each s10 variant, we plotted individual relative emission spectra (**Figure 4A**) relative to WT. We observed no fluorescent complementation when LOO10-GFP was incubated with the L207R variant; the spectrum of this complex was unchanged relative to LOO10-GFP. S10 WT complemented LOO10-GFP with 5-fold increase in 520 nm signal, but no additional effect on brightness when introducing H199Y as a single substitution. T203Y variant caused a spectral shift of the emission maximum of the recovered FP complex from 506 to 520 nm, offering ~ 1.6-fold higher brightness of the T203Y variant compared to WT at 520 nm. H199Y in combination with T203Y increased the 520 emission by 1.8-fold compared to WT. The loss-of-function effect of L207R and the gain-of-function effects of T203Y variants were both captured by the microarray fluorescence of these variants. Still, brightness effects in solution could not explain the microarray data; a R^2^ < 0.5 being obtained when correlating the two datasets (**Figure 4B**). In particular, H199Y, with a 10-fold increase in microarray fluorescence as a single substitution, showed no brightness effects when saturating LOO10-GFP with this variant in solution.

**Figure 4.**
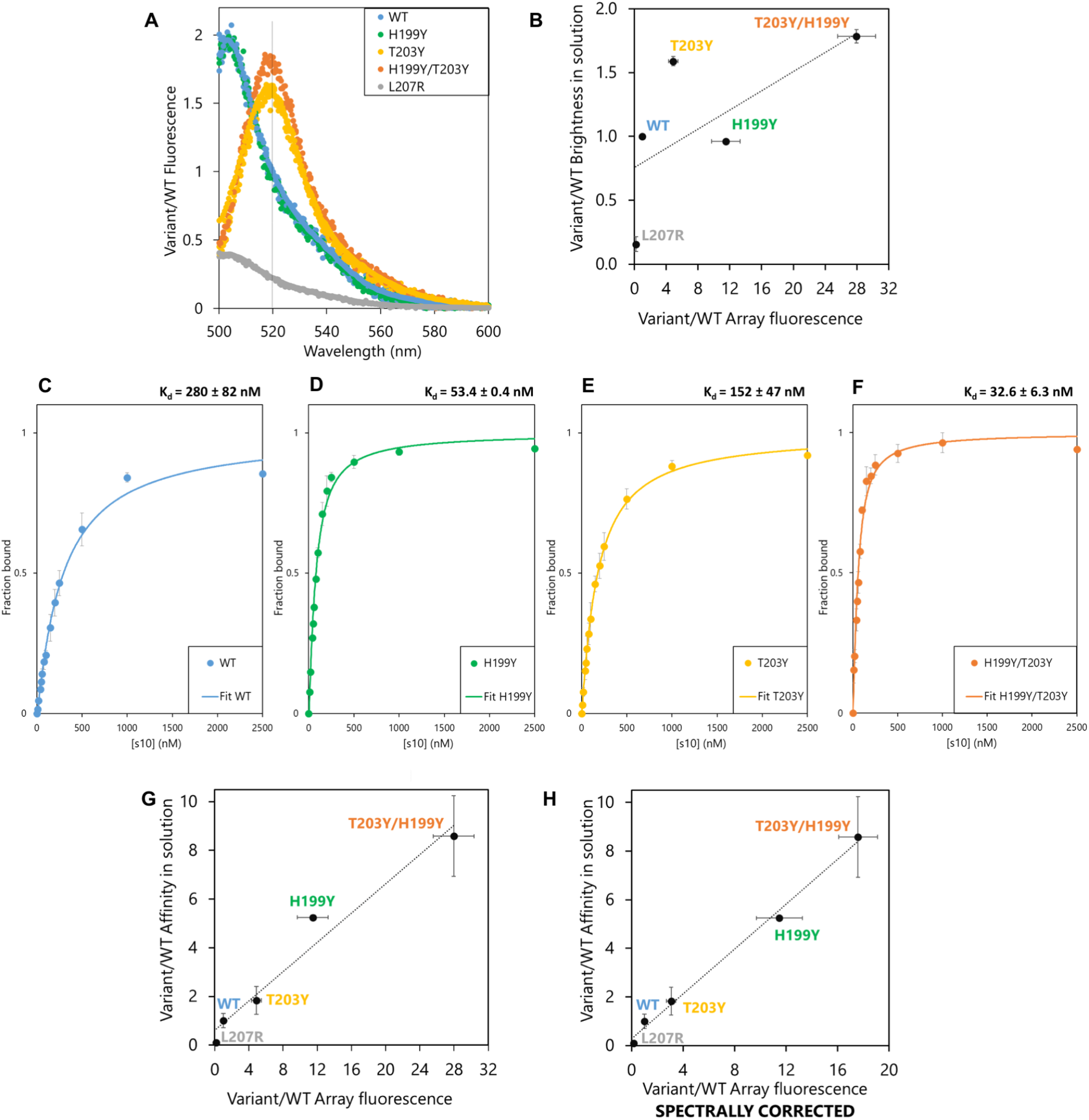
Brightness and affinity of selected s10 substitutions in solution. (A) Fluorescence emission spectra of LOO10-GFP saturated with each of the 5 s10 variant peptides and excited at 488 nm, normalized to emission of WT complex at 520 nm (marked on the plot with a dashed line). Spectra of 3 independent replicas for each peptide variant are overlaid in the plot. (B) Correlation between relative microarray fluorescence and relative brightness of the FP complexes in solution. (C-F) Binding isotherms of (C) WT, (D) H199Y (390ex/506em) and (E) T203Y, (F) H199Y/T203Y (495ex/520em) variants. Error bars represent standard deviation of 3 independent measurements. (G) Correlation between relative microarray fluorescence and relative affinity of the tested split fragments in solution agrees well with a linear model. (H) Combining the spectral and affinity properties in solution still explains the microarray fluorescence.

Next, we titrated LOO10-GFP with increasing concentrations of s10 variant in solution and fitted the binding curves (**Figure 4C-F**) to calculate dissociation constants, *K*_d_, as a measure of affinity between the fragments. The H199Y substitution showed a ~ 5-fold increase in affinity compared to WT, while the double substituted H199Y/T203Y had a ~ 9-fold increase. The ~ 2-fold increase in affinity of T203Y relative to WT previously reported for a 19-mer s10 (12) is replicated by our data. Competition experiments suggested that L207R variant binds LOO10-GFP very weakly and is almost fully displaced by a 4-fold lower concentration of WT variant (**Figure S7**). We therefore approximated the affinity of L207R to be at least 10-fold weaker than WT.

Assuming a sub-saturation regime where *K*_d_ is greater than the LOO10-GFP concentration, the association constants, *K*_d_^-1^, should scale linearly with array fluorescence, see Methods. Under this assumption, affinity effects are more likely to explain the array fluorescence, although they do not take into account the spectral contributions (**Figure 4G**). Furthermore, taking the spectral shift for T203Y and H199Y/T203Y into account, the spectrally corrected microarray fluorescence offered a slightly better fit, although probably not significant (**Figure 4H**). Based on these experiments, we conclude that with affinities in the nM range, the signal intensity on peptide microarrays in this format faithfully reflects the binding affinity between the split fragments.

## DISCUSSION

We have developed a precise, robust and accurate method for exploring the substitutional landscape for leave-one-out split fluorescent proteins, that has generated a comprehensive sequence-function map of a splitFP tag. Chemical synthesis of peptides in the library avoided time-consuming and bias-inducing steps like cloning, expression, purification or sequencing, that are usually required in genetic screens. By having full control over the s10 sequence on the microarray, we circumvented the limitations of the DNA codon table. Because double and triple substitutions in a given codon are rare, random mutagenesis will typically only generate a subset of mutations (21) and will never exhaustively sample double or triple amino acid substitutions. In the microarray setup it is possible to investigate every peptide in the library independently of Hamming distance from the starting sequence at the DNA level. Thus, libraries can be set up highly diverse (potentially including non-natural amino acid residues) offering a one-to-one picture of the entire functional/binding landscape, including low and medium performing sequences. In addition, because the LOO-FP chromophore is matured prior to our assay, we accessed high affinity s10 variants that might not be discoverable by multiplexed expression of the full-length proteins, where the chromophore cyclisation and oxidation is a prerequisite for fluorescence. Indeed, the H199Y substitution either does not show up or is identified as likely destabilizing in other GFP genetic screens (21, 22). It is an interesting possibility that this variant, while stabilizing the mature GFP, is disfavored in genetic screens because it does not stabilize the non-fluorescent immature precursor. Lastly, our assay proposed a direct measure of peptide binding in high-throughput, avoiding false positives caused by oligomeric fluorescent species that can appear in genetic selection experiments performed in cells (9).

We should point out that the interpretation of results in the microarray platform assumes a similar chemical yield of peptide across variants. The level of reproducibility between 12 replicas suggested intra-sequence synthesis yields are very similar. Inter-sequence yields were more difficult to assess, but the consistently lower signals when incorporating histidine in any part of s10 might suggest a reduced coupling yield when histidine is incorporated in the sequence. However, low histidine coupling yield was not observed in another study using microarrays from the same manufacturer (14).

In this analysis, we identified several interesting GFP strand 10 peptide substitutions and truncations. In particular, we note a double substitution, H199Y/T203F, which presented 54-fold higher microarray fluorescence compared to the WT sequence. We also found that the H199Y/T203Y s10 variant had almost 10-fold higher solution affinity compared to WT. Truncations of s10 to 11-mer or even 9-mer proved active on the peptide microarray and the 11-mer also effectively reconstituted fluorescence as a free peptide in solution. These short versions of s10 with the H199Y substitution could be readily used for *in vitro* applications, and could possibly be further investigated as non-interfering protein tags due to their small size.

For developing this screening platform, we used the split 10 FP system as a model, mainly due to its high affinity *in vitro*. Still, the LOO10-GFP efficiency *in cellulo* proved poor in previous studies (6, 23). For engineering improved *in cellulo* and *in vivo* tags using the microarray platform reported here, one could turn to the strand 7 and strand 11 LOO systems (23). By incubating a microarray library with an immature LOO-FP, one could study what the sequence requirements would be for both binding and chromophore maturation. Indeed, β-strand-assisted chromophore maturation of LOO11-GFP on solid support has already been demonstrated (24).

Reaching saturation for every peptide could theoretically offer the possibility of detecting variants with intrinsic FP brightness improvement, desirable for many applications. A limitation in our study was that LOO10-GFP is a rather unstable protein with fairly low solubility, 2 μM being the maximum reliable concentration we could obtain for our microarray experiments. A more stable LOO10-GFP might be desirable, since titrating the microarray with increasing concentrations of LOO10-GFP could saturate all peptides and possibly allow plotting full binding curves for each peptide. This possibility was demonstrated in previous microarray screens (25, 26). In the absence of strand 10, on the other hand, a hydrophobic patch is exposed, thus mutations that stabilize LOO10-GFP, *e.g*. by making this surface more hydrophilic, would also likely decrease affinity to strand 10.

We believe the platform reported here can be generalized to many split fluorescent and luminescent proteins. Screening on microarrays may be limited by unspecific binding which is avoided for split systems that require specific complementation in order to function. Different color FPs could be studied, since microarray scanners employ excitation laser sources emitting in the blue, green and red spectrum regions and provide multiple emission filter options. Tuning protein concentrations to the sub-saturation regime is typically not difficult, however, requires a bright chromophore and a sensitive microarray reader. For reasons of sensitivity, assaying split proteins using this technology is most likely limited to fluorescence detection. Thus, studying split enzymes systems would likely require appropriate (insoluble) fluorescent product formation. One possibility is using internally quenched substrates, similar to those used in some protease assays (27).

Overall, full control over the desired substitutions on peptide microarrays makes them versatile alternatives to cell-based approaches for rational design of binders, massively parallel testing of computational design and benchmarking biophysical prediction methods.

## METHODS

### Preparation of LOO10-GFP

We used an adapted method from (12). Engineered plasmids of full-length circularly permuted superfolder GFP in pET-15b vectors were kindly provided by Steven Boxer, Stanford University. The full-length GFP was expressed in BL21 (DE3) cells in commonly used AB-LB growth media, where ampicillin (VWR) was added to a concentration of 100 μg/ml and glucose to a concentration of 1% (w/v). The starter culture was grown overnight at 37°C in LB medium, then expanded in AB-LB medium in a ratio of 1:100 starter culture to growth medium. This culture was grown at 37°C until an OD600 of ~0.6, induced with 1 mM isopropyl β-d-1-thiogalactopyranoside (Sigma) and agitated at 17°C overnight for protein expression. The cells were harvested on a Lynx 4000 (SORVALL) centrifuge at 20,000g for 30 minutes. The cell pellet was resuspended in lysis buffer 50 mM Hepes, 300 mM NaCl, pH 8 at 25 ml buffer per liter culture and sonicated in an UP2009 (Hielscher) for 8 cycles 30 sec pulse / 30 sec pause. After one round of 13,000g centrifugation for 15 minutes, the supernatant was poured onto a Ni-NTA column (GE Healthcare) equilibrated with lysis buffer. Three column volumes of lysis buffer supplemented with 20 mM imidazole (Merck) was used for washing, and the same buffer supplemented 200 mM imidazole was used to elute the protein. Imidazole was removed by dialyzing the sample through a Spectra/Por 3.5 kDa membrane into lysis buffer overnight at 4°C.

The loop between strand 10 and 11 was digested using a 0.5 μM trypsin (DIFCO) solution. The reaction contained full length circularly permuted GFP and 1% (M/M) trypsin in cleavage buffer 50 mM Tris, 20 mM CaCl2 (Merck), pH 8. The tube was incubated for 30 minutes at room temperature, before the trypsin was inhibited with protease inhibitor phenylmethylsulfonyl fluoride (Sigma), at a final concentration of 1 mM. The mix was then precipitated in a 60% ammonium sulfate solution and centrifuged at 5000g for 10 minutes at 4°C. The pellet was dissolved in denaturation buffer (50 mM Hepes, 50 mM NaCl, 6 M GdnHCl, pH 8) and loaded on a Superdex 75 10/300 GL (GE Healthcare) size exclusion column to remove s10. LOO10-GFP was eluted isocratically in denaturation buffer at 0.6 ml/min. Fractions with absorption A447 > 0.1 (1 cm path length) were pooled together and stored at 4°C until usage. For quantitative measurements and microarray experiments, LOO10-GFP was refolded by desalting on an Illustra NAP5 column (GE Healthcare) equilibrated with assay buffer 50 mM Hepes, 100 mM NaCl, 0.1% v/v Tween20, pH 8 using the manufacturer’s protocol.

### Synthetic peptides

Chemically synthesized s10 peptides NHYLSTQTVLS (WT), NYYLSTQTVLS (H199Y), NHYLSYQTVLS (T203Y), NYYLSYQTVLS (H199Y/T203Y), LPDNHYLSTQTVRSKDPNE (L207R) were purchased from Schafer-N or TAG Copenhagen with >95% purity.

### Fluorescence spectroscopy

Spectral measurements were performed by mixing 30 nM LOO10-GFP with 3 μM (100 fold excess) of each s10 peptide variant (WT, L207R, H199Y, T203Y or H199Y/T203Y) and incubated ~ 1 hour in assay buffer for full complementation. Fluorescence emission spectra were recorded by exciting samples at 488 nm (5 nm slit) and a 5 nm emission slit in the 495-600 nm range on a PerkinElmer LS 55 Luminescence Spectrometer at 25°C. Spectra were taken in 3 independent replicas for each peptide variant.

Affinity measurements were performed by mixing 50 nM LOO10-GFP with solutions of increasing concentrations of each s10 peptide variant WT, H199Y, T203Y or H199Y/T203Y (0, 10, 20, 40, 50, 60, 80, 100, 150, 200, 250, 500, 1000, 2500 nM) and incubating overnight at 4°C for equilibration. Fluorescence of each sample was measured at 390/506 nm for T203 variants and at 495/520 nm for Y203 variants, at 25°C, excitation slit 12 nm, emission slit 20 nm and 10s integration time. The affinity experiments were performed in 3 independent replicas for each peptide variant. Fluorescence data for each curve were fit to a quadratic equation:

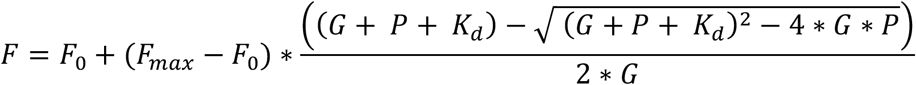

where F_0_ was background fluorescence (FU), F_max_ was fluorescence at saturation (FU), G was LOO10-GFP concentration (nM), P is s10 peptide concentration (nM) and *K*_d_ is the binding affinity between the two fragments (nM). All fits were performed in OriginPro 2017.

### Absorption spectroscopy

Concentration of LOO10-GFP was estimated by measuring absorbance at 447 nm and using its extinction coefficient at 0.1 M NaOH of Ɛ_447_ = 44,100 M^-1^ cm^-1^ (12). The s10 peptides were dissolved in assay buffer and the predicted Ɛ_280_ based on tryptophan and tyrosine content was used for concentration determination (28). All absorbance measurements were done on a Perkin Elmer Lambda 35 Spectrophotometer.

### Microarray incubation and data analysis

The peptide library was synthesized on a single chip with 12 identical sectors, each containing 15,301 peptide spots of 20 x 20 μm, 183,612 peptide spots in total. Out of these, 12,544 peptide variants x 12 = 150,528 spots were analyzed for this report. The remaining 2757 spots were part of another project and are not reported here.

Before incubation with LOO10-GFP, the array surface was hydrated with assay buffer for 10 minutes at 25°C under shaking. After removal of the buffer, 4 mL solution of 2 μM LOO10-GFP in assay buffer was added to the array and incubated over night (~ 18 hours) at 4°C under shaking. The array was washed two times in assay buffer at room temperature and dried under a gentle stream of N2. The array was imaged on an InnoScan 1100 AL (INNOPSYS) microarray laser scanner at 488 nm excitation and with a 520±5 nm emission bandpass filter. Images were collected at low laser power (5 mW) with gain of 20, at 1 μm resolution. Fluorescence data were extracted from images using ImageJ MicroArray Profile plugin Version 3.1. with rectangle ROIs of 20 x 20 μm and assigned to each peptide variant spot.

Any outliers caused by dust or other contaminations were estimated by observing the distribution of the 12 replicas for each peptide. We chose a conservative cutoff for outlier removal, since contaminants should be very bright compared to a regular high signal. Thus, all the values within 6 fold Median Absolute Deviation (MAD) from the median for each peptide were taken into further analysis, while the rest were eliminated as outliers. MAD is generally not influenced by outliers, so the observation that the standard deviation after outlier removal is similar to MAD before outlier removal (**Figure S8)** demonstrates that most outliers were successfully eliminated.

As each of the 12 sectors were, in principle, identical, the average sector signal should be the same. However, the fact that average brightness shifted in a consistent gradient across the slide suggested an artifactual global variation (**Figure S9**). This could be due to inhomogeneity of the functional surface or inhomogeneity of buffer component allocation during incubation/washing steps. However, when plotting all fluorescent signals of each sector against the others, we observed that the variant libraries in all 12 sector replica correlated well with each other, with Pearson coefficients > 0.94 (**Figure S1**), therefore we considered appropriate normalizing the mean fluorescence of each sector to the mean fluorescence of the microarray. When examining the impact of normalization on data quality, we observed that the mean variant fluorescence of 12 replicas was maintained constant after normalization (**Figure S10A**), while the standard deviation values dropped ~ 2 fold (**Figure S10B**). Thus, the normalization procedure did not quantitatively change the fluorescence signals of variants, but only made the data more precise. The array background was removed for each peptide by subtracting the mean fluorescence of blank spots in each sector. All negative/zero fluorescence values after background removal were replaced with the value +1, to avoid issues at log2 transformation.

We measured the mean fluorescence and standard deviation for the s10_long_ WT peptide at 741 ± 270 fluorescence units (FU) for non-normalized data and 741 ± 64 FU for normalized data. Within the same peptide format, we considered as “WT-like fluorescence” all variant means within 2 standard deviations (~ 95% confidence interval) from the non-normalized WT mean, corresponding to ~ 4.3 standard deviations (> 99.99% confidence interval) from the normalized WT mean. Variant fluorescence means above the WT interval were defined as gain-of-function or enhancing, while variant means below the WT interval were defined as loss-of-function or detrimental throughout the paper.

Substitutional tolerance at each s10 position was calculated by counting the number of substitutions in that position which cause a change in fluorescence, either gain-of-function or loss-of-function (as defined previously). We subtracted this number from the total 19 substitutions at each position.

If not stated otherwise, all variant fluorescence values are outlier-and-background-removed normalized mean variant fluorescence of 12 replica. All plots are generated using the absolute mean variant fluorescence, except all heatmaps and variant effect plots where we used log2(mean variant fluorescence), and the experimental validation dataset where we used relative variant fluorescence against WT *i.e*. mean variant fluorescence/mean WT fluorescence.

### Thermodynamic description of microarray binding and spectral correction

The equilibrium between LOO10-GFP, *P*, immobile s10 peptide, *S*, and the fluorescent complex, *PS*, gives the fraction of fluorescent chromophores per microarray spot:

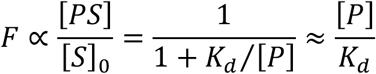

We assume this to be proportional to the observed fluorescence, *F*, and that all spots have the same amount of peptide |*S*|_0_ = [*S*] + [*PS*]. The last approximation describes the sub-saturation regime > [*P*].

For spectral correction, the microarray fluorescence of T203Y and H199Y/T203Y is simply divided by the spectral shift factor of 1.6 determined from the Figure 4A.

## Supporting information

Sup_Info_legend_Txt

## Data and code availability

The raw data from the microarray screen and the R code used to clean, sort and plot the data are available at https://github.com/onea7/Substitutional-landscape-splitFP.

## ACKNOWLEDGEMENTS

We wish to thank Steven Boxer (Stanford University) for help with the cp-sfGFP plasmids and suggestions; Claus Schafer-Nielsen (Schafer-N A/S) for many discussions on the microarray layout and handling; Julie Eilskov Bolding and Selma Kofoed Bendtsen (University of Copenhagen) for performing preliminary experiments that helped us better understand the splitFP system. This work was supported by Independent Research Fund Denmark.

## AUTHOR CONTRIBUTIONS

O.N.A. and J.R.W. conceived the project and designed the experiments; O.N.A., A.R., N.A.M.D., R.P. and A.N-M. performed the experiments; O.N.A., K.E.J. and D.F.H. analyzed the data; O.N.A. and J.R.W. wrote the manuscript with input from all authors.

## COMPETING INTERESTS

The authors declare no competing interests.

## Notes

https://github.com/onea7/Substitutional-landscape-splitFP

### Competing Interest Statement

The authors have declared no competing interest.

## REFERENCES

1. Ghosh I, Hamilton AD, Regan L. Antiparallel leucine zipper-directed protein reassembly: Application to the green fluorescent protein. Journal of the American Chemical Society. 2000;122(23):5658–9.

2. Cabantous S, Terwilliger TC, Waldo GS. Protein tagging and detection with engineered self-assembling fragments of green fluorescent protein. Nat Biotechnol. 2005;23(1):102–7.

3. Cabantous S, Nguyen HB, Pedelacq JD, Koraichi F, Chaudhary A, Ganguly K, et al. A New Protein-Protein Interaction Sensor Based on Tripartite Split-GFP Association. Sci Rep-Uk. 2013;3:2854.

4. Romei MG, Boxer SG. Split Green Fluorescent Proteins: Scope, Limitations, and Outlook. Annu Rev Biophys. 2019;48:19–44.

5. Pedelacq JD, Cabantous S. Development and Applications of Superfolder and Split Fluorescent Protein Detection Systems in Biology. Int J Mol Sci. 2019;20(14):3479.

6. Huang YM, Nayak S, Bystroff C. Quantitative in vivo solubility and reconstitution of truncated circular permutants of green fluorescent protein. Protein Sci. 2011;20(11):1775–80.

7. Feng SY, Sekine S, Pessino V, Li H, Leonetti MD, Huang B. Improved split fluorescent proteins for endogenous protein labeling. Nat Commun. 2017;8:370.

8. Kamiyama D, Sekine S, Barsi-Rhyne B, Hu J, Chen BH, Gilbert LA, et al. Versatile protein tagging in cells with split fluorescent protein. Nat Commun. 2016;7:11046.

9. Huang YM, Banerjee S, Crone DE, Schenkelberg CD, Pitman DJ, Buck PM, et al. Toward Computationally Designed Self-Reporting Biosensors Using Leave-One-Out Green Fluorescent Protein. Biochemistry-Us. 2015;54(40):6263–73.

10. Blanco C, Janzen E, Pressman A, Saha R, Chen IA. Molecular Fitness Landscapes from High-Coverage Sequence Profiling. Annu Rev Biophys. 2019;48:1–18.

11. Szymczak LC, Kuo HY, Mrksich M. Peptide Arrays: Development and Application. Analytical Chemistry. 2018;90(1):266–82.

12. Do K, Boxer SG. Thermodynamics, kinetics, and photochemistry of beta-strand association and dissociation in a split-GFP system. J Am Chem Soc. 2011;133(45):18078–81.

13. Dixon AS, Schwinn MK, Hall MP, Zimmerman K, Otto P, Lubben TH, et al. NanoLuc Complementation Reporter Optimized for Accurate Measurement of Protein Interactions in Cells. ACS Chem Biol. 2016;11(2):400–8.

14. Buus S, Rockberg J, Forsstrom B, Nilsson P, Uhlen M, Schafer-Nielsen C. High-resolution mapping of linear antibody epitopes using ultrahigh-density peptide microarrays. Mol Cell Proteomics. 2012;11(12):1790–800.

15. Wang W, Woodbury NW. Selective protein-peptide interactions at surfaces. Acta Biomater. 2014;10(2):761–8.

16. Wang W, Woodbury NW. Unstructured interactions between peptides and proteins: exploring the role of sequence motifs in affinity and specificity. Acta Biomater. 2015;11:88–95.

17. Tsien RY. The green fluorescent protein. Annu Rev Biochem. 1998;67:509–44.

18. Jung G, Wiehler J, Zumbusch A. The photophysics of green fluorescent protein: Influence of the key amino acids at positions 65, 203, and 222. Biophys J. 2005;88(3):1932–47.

19. Heikal AA, Hess ST, Baird GS, Tsien RY, Webb WW. Molecular spectroscopy and dynamics of intrinsically fluorescent proteins: coral red (dsRed) and yellow (Citrine). Proc Natl Acad Sci U S A. 2000;97(22):11996–2001.

20. Fujiwara K, Toda H, Ikeguchi M. Dependence of alpha-helical and beta-sheet amino acid propensities on the overall protein fold type. BMC Struct Biol. 2012;12:18.

21. Lars Behrendt AS, Shiraz Ali Shah, Karsten Zengler, Søren J. Sørensen, Kresten Lindorff-Larsen, Jakob R. Winther. Deep mutational scanning by FACS-sorting of encapsulated E. coli micro-colonies. BioRxiv. 2018.

22. Sarkisyan KS, Bolotin DA, Meer MV, Usmanova DR, Mishin AS, Sharonov GV, et al. Local fitness landscape of the green fluorescent protein. Nature. 2016;533(7603):397–401.

23. Huang YM, Bystroff C. Complementation and reconstitution of fluorescence from circularly permuted and truncated green fluorescent protein. Biochemistry-Us. 2009;48(5):929–40.

24. Lundqvist M, Thalen N, Volk AL, Hansen HG, von Otter E, Nygren PA, et al. Chromophore pre-maturation for improved speed and sensitivity of split-GFP monitoring of protein secretion. Sci Rep-Uk. 2019;9.

25. Layton CJ, McMahon PL, Greenleaf WJ. Large-Scale, Quantitative Protein Assays on a High-Throughput DNA Sequencing Chip. Mol Cell. 2019;73(5):1075–82 e4.

26. Katilius E, Flores C, Woodbury NW. Exploring the sequence space of a DNA aptamer using microarrays. Nucleic Acids Res. 2007;35(22):7626–35.

27. Poreba M, Szalek A, Rut W, Kasperkiewicz P, Rutkowska-Wlodarczyk I, Snipas SJ, et al. Highly sensitive and adaptable fluorescence-quenched pair discloses the substrate specificity profiles in diverse protease families. Sci Rep. 2017;7:43135.

28. Pace CN, Vajdos F, Fee L, Grimsley G, Gray T. How to measure and predict the molar absorption coefficient of a protein. Protein Sci. 1995;4(11):2411–23.

