## Supplementary material for "Substitutional landscape of a split fluorescent protein fragment using high-density peptide microarrays": Sup_Info_legend_Txt

SUPPLEMENTARY INFORMATION
Substitutional landscape of a split fluorescent protein fragment
Oana N. Antonescu-1), Andreas Rasmussen-1), Nicole A.M. Damm-1), Ditte F. Heidemann-1), Roman Popov-2), Alexander Nesterov-Mueller-2), Kristoffer E. Johansson-1) & Jakob R. Winther-1)
1)Linderstrøm-Lang Centre for Protein Science, Section for Biomolecular Sciences, Department of Biology, University for Copenhagen, Ole Maaloes Vej 5, DK-2200 Copenhagen N, Denmark
2)Institute of Microstructure Technology, Karlsruhe Institute of Technology, Hermann-von-Helmholtz-Platz 1, 76344, Eggenstein-Leopoldshafen, Germany


Table S1. Overview of the composition of the peptide library. Strand 10 substitutional variants and controls synthesized on the microarray. The linker sequences are gs2: GS; gs7: GSGSGSG, gk7: GKGSKSG and ge7: GEGSESG.

Figure S1 . Correlations between the 12 sectors. Correlation of variant fluorescence between all 12 sector replicas, after outlier removal but before sector normalization. The Pearson correlation coefficients are indicated in the upper matrix.

Figure S2. Linker effects in the short format.  Correlations between signals from substitutions in s10short with different linkers: no linker, positive GKGSKSG (gk7) linker, negative GEGSESG (ge7) linker, amd neutral GSGSGSG (gs7) linker. The Pearson correlation coefficients between the different substitutional libraries are indicated in the upper matrix.

Figure S3. Left- and right-shifting effects. s10short versus s10shortL and s10shortR. All substitution variants of the short format with ge7 (red), gk7 (green), gs7 (dark blue) and no linker (light blue) on the x-axes compared to the same substitution and linker variants of the (A) shortL and (B) shortR formats on the y-axis. N- and C-terminus variants in the shifted formats that are not found in the short format are marked with magenta (NT) or grey (CT) crosses.

Figure S4. Effect of context on the fluorescence of substitutional variants. Comparing single (WT context), double (H199Y, T203Y, T203I, L207V and L207R contexts) and triple (H199Y/T203Y context) substitutions from s10long WT. In each context, the jitter points are colored based on the position substituted, resulting in 20 points of the same color per position. Each variant is presented as mean over 12 replica ± one standard deviation as error bar. The dashed lines delimit the “WT interval” i.e. mean WT fluorescence ± 4.3 standard deviations. The interval above WT is considered GoF (gain-of-function), while the interval below LoF (loss-of-function).

Figure S5. Top fluorescing variants. (A) Box plot of top 9 highest fluorescing variants of the long format, representing the core sequence and flanking regions. (B) Box plot of top 5 highest fluorescing variants the short-gk7 format, representing only the core sequence. In both (A) and (B), each box indicates the distribution of absolute fluorescence of each variant across the 12 replica. Box plots of some single substitutions and the H199Y/T203Y double substitution are added for comparison.

Figure S6. Proposed model for interaction at the peptide microarray surface. Higher microarray fluorescence at peptide field A/C than at peptide field B/D can occur due to: (left) Sub-saturation conditions: Higher affinity of peptide A than peptide B to LOO10-GFP, resulting in more complemented FP molecules on peptide field A than B or (right) Saturation conditions: Higher brightness of FP complex C than D, at equal number of complemented FP molecules on peptide field C and D.

Figure S7. Competition studies with L207R variant. Time resolved fluorescence of association between 100 nM LOO10-GFP and 2.5 µM s10short WT, with – blue curve - or without – red curve – pre-incubation with 10 µM s10long L207R. 

Figure S8. Quality control of the microarray data after outlier removal. (A) Scatter plot comparing the median absolute deviation (MAD) across 12-replica for each variant before outlier removal and the standard deviation (SD) across 12-replica for the same variant after outlier removal.

Figure S9. Fluorescence across the 12 microarray sectors. Heatmap of mean fluorescence per sector, before normalization. Color key: Linear gradient between grey for dim sectors and green for bright sectors.

Figure S10. Quality control of the microarray data after cleaning. (A) Scatter plot comparing the median absolute deviation (MAD) across 12-replica for each variant before outlier removal and the standard deviation (SD) across 12-replica for the same variant after outlier removal. (B) Mean and (C) SD across 12-replica for each variant before normalization plotted against (B) mean and (C) SD of the same variant after normalization.
