## Supplementary figures and images for "Substitutional landscape of a split fluorescent protein fragment using high-density peptide microarrays"

### Sup_Info_Figure_S1

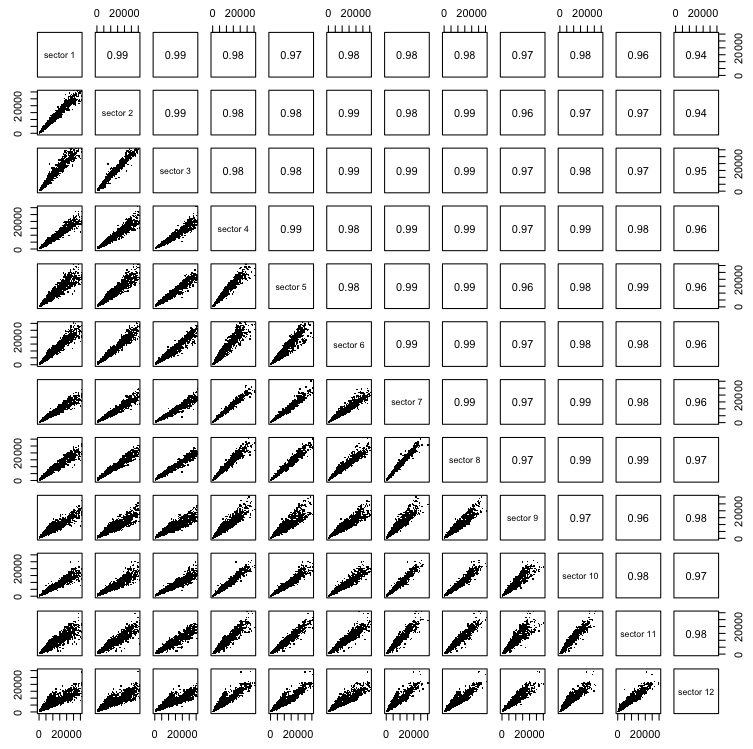

### Sup_Info_Figure_S2

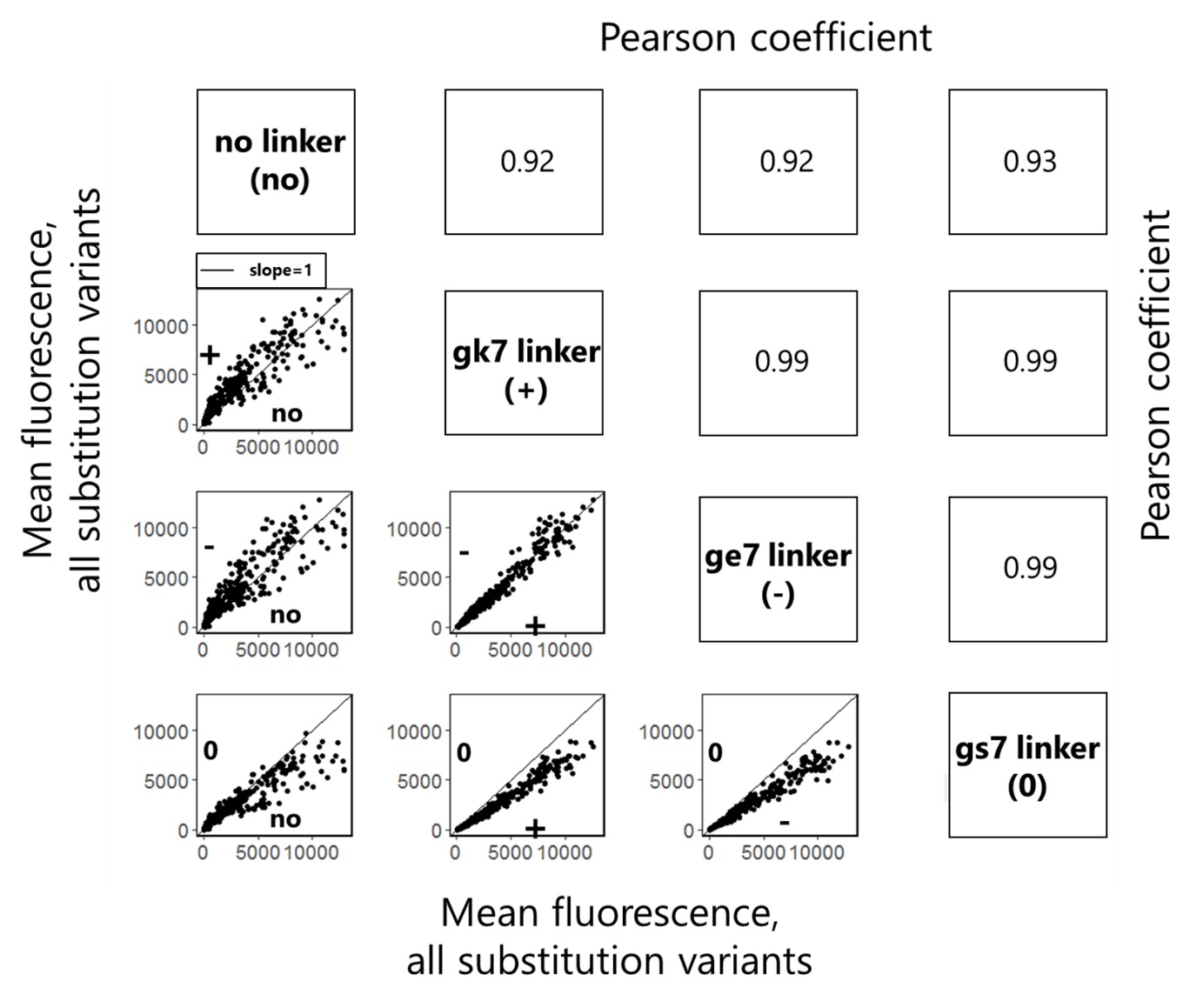

### Sup_Info_Figure_S3

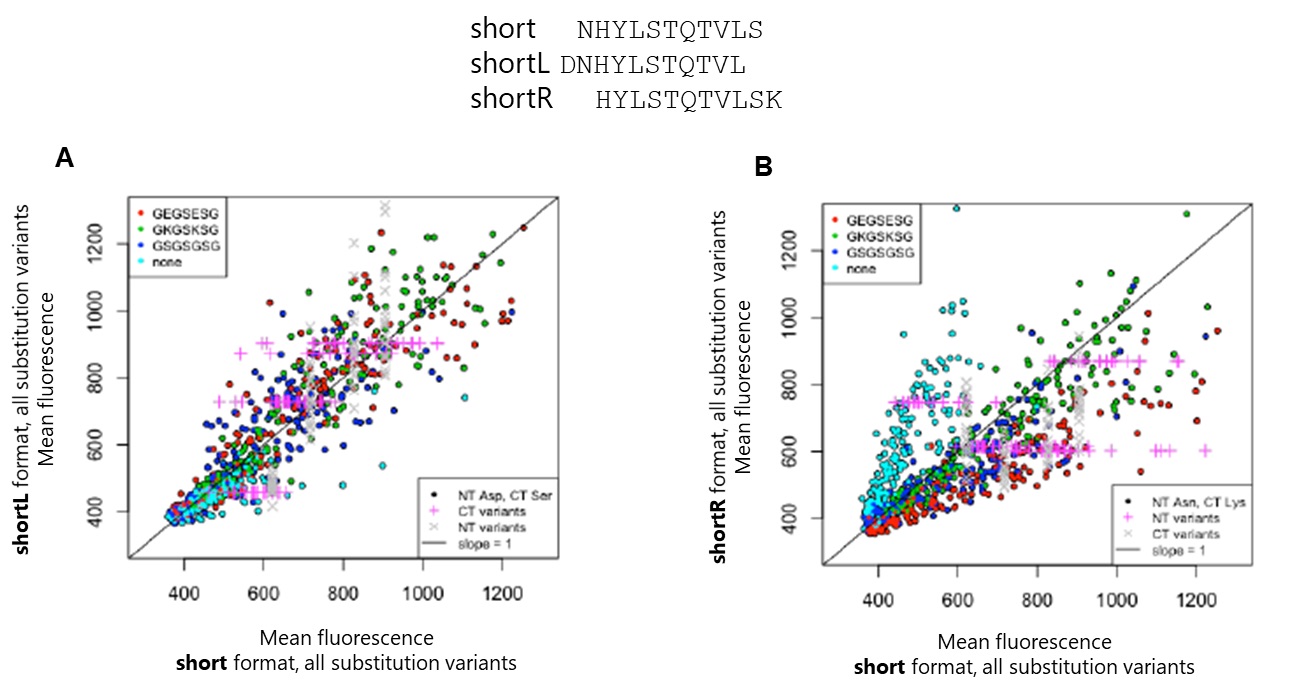

### Sup_Info_Figure_S4

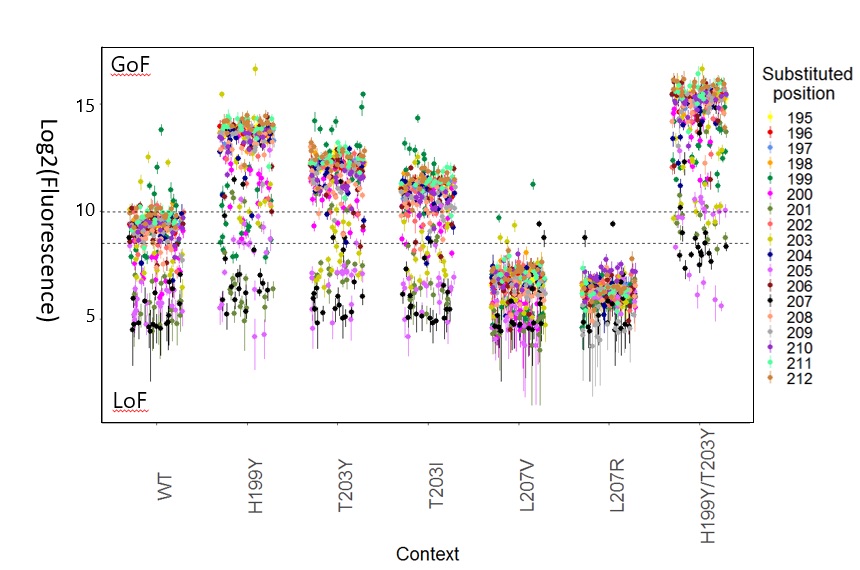

### Sup_Info_Figure_S5

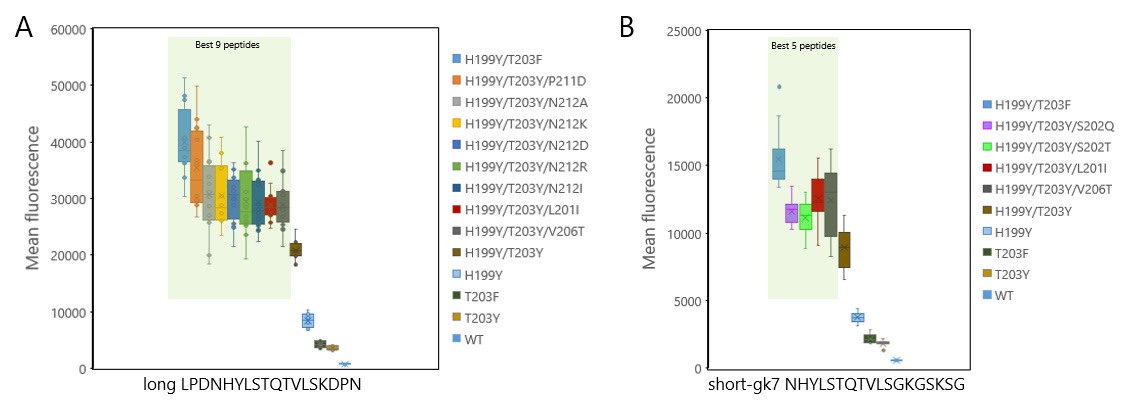

### Sup_Info_Figure_S6

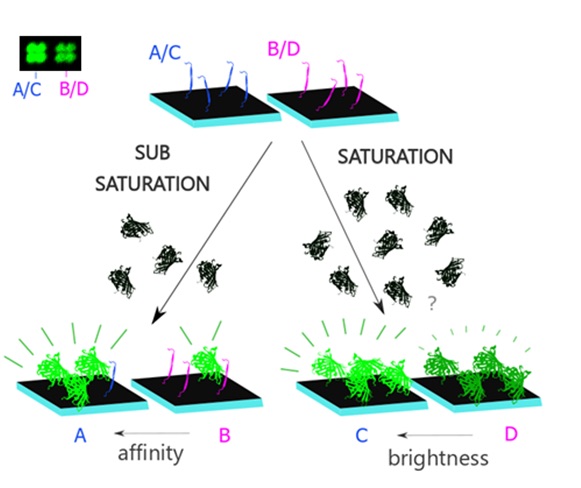

### Sup_Info_Figure_S7

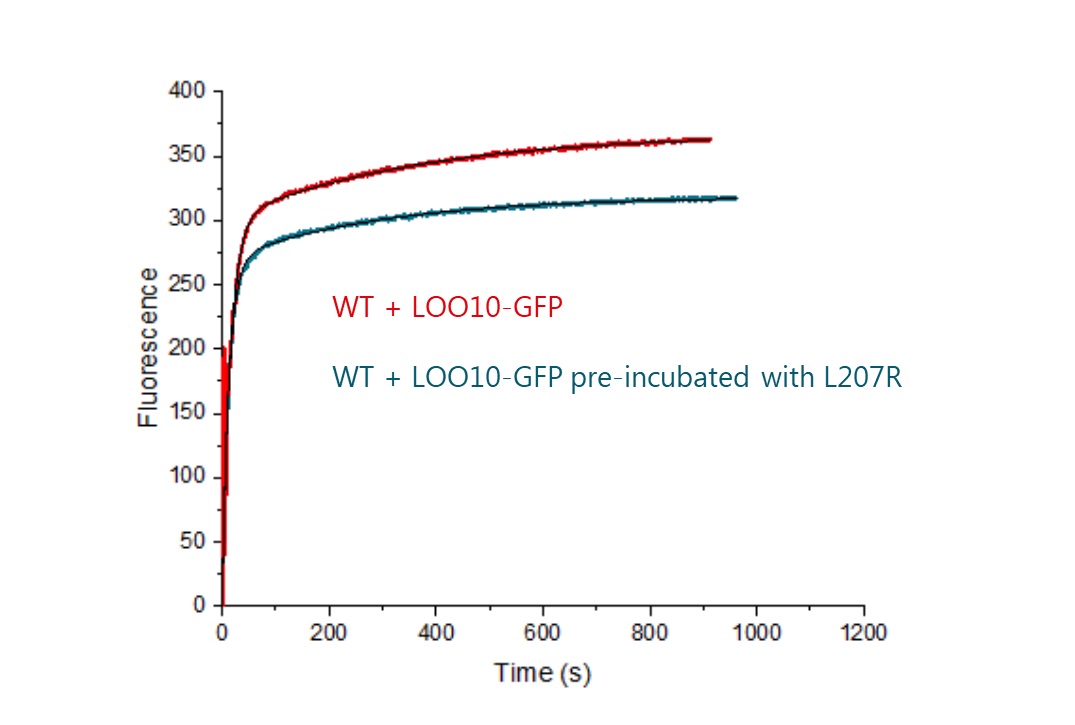

### Sup_Info_Figure_S8

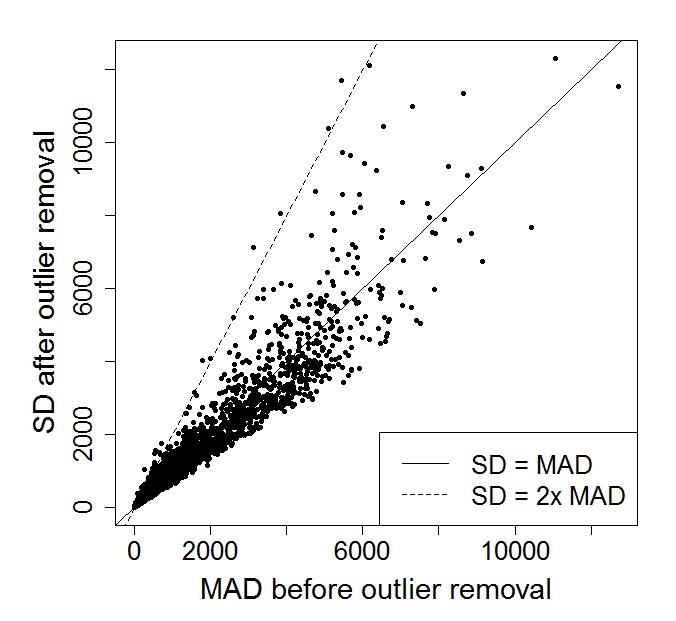

### Sup_Info_Figure_S9

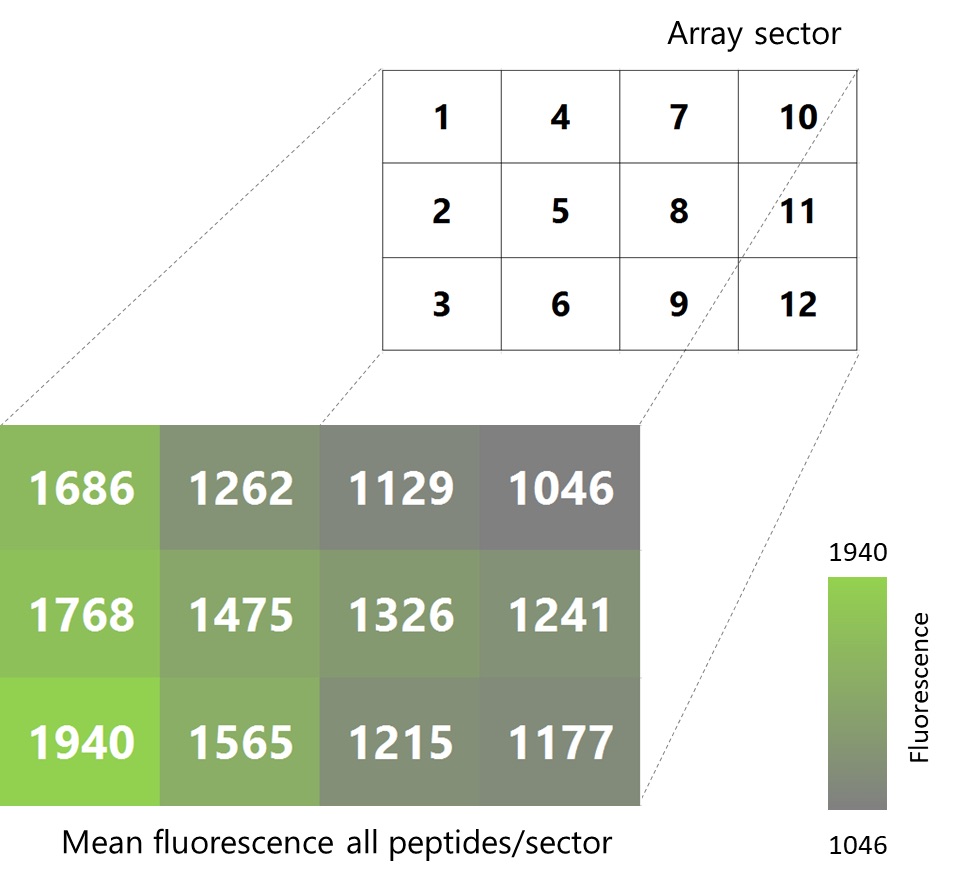

### Sup_Info_Figure_S10

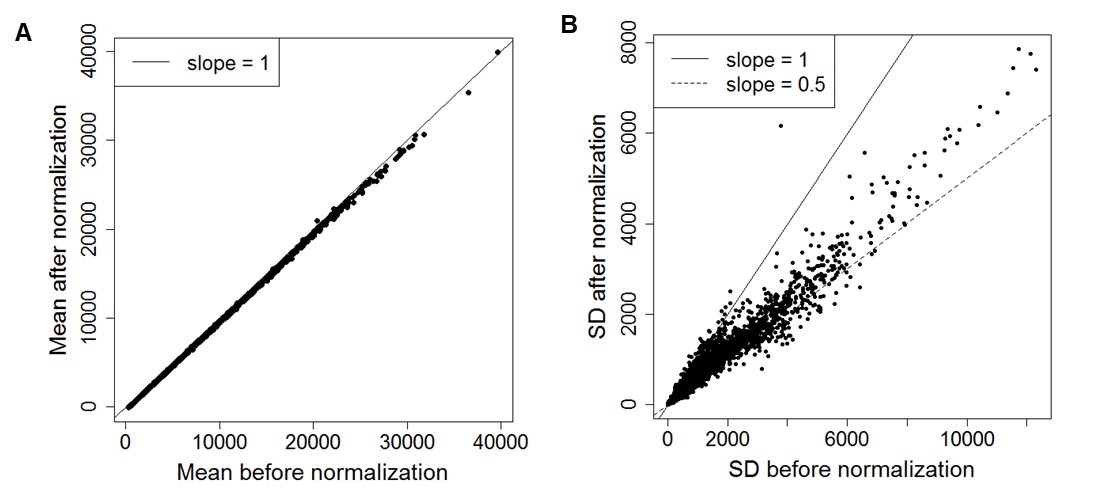

### Sup_Info_Table_S1

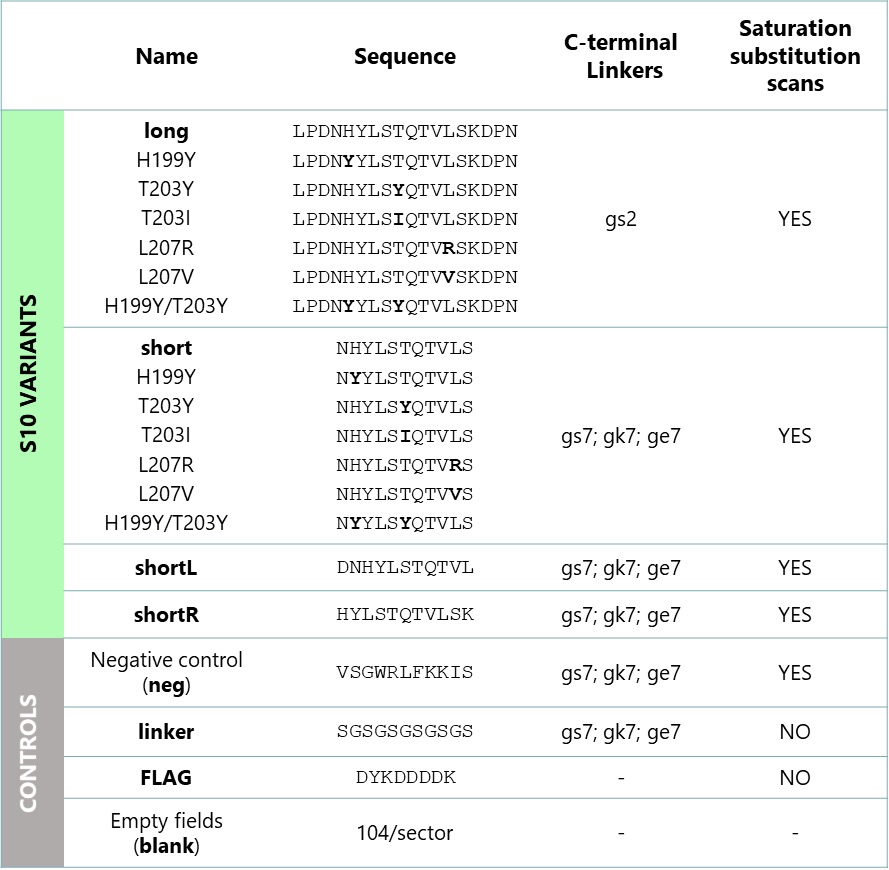
